# Injury-induced Cxcl11 and neutrophil signaling drive zebrafish kidney regeneration by generating a nephrogenic niche of Fgf and Wnt expression

**DOI:** 10.64898/2026.04.08.717325

**Authors:** Olaleye Olajuyin, Heiko Schenk, Will G.B. Sampson, Omodasola Adekeye, Caramai N. Kamei, Rohan M. Upadhyay, Ryan Kennedy, Emma Morrison, Riley Callahan, Frederic Bonnet, Joel Graber, Ryan Seaman, Heath Fuqua, Robert Wheeler, Leif Oxburgh, Iain A. Drummond

## Abstract

Adult zebrafish regenerate their kidneys after injury by activating quiescent renal stem cells, however the injury signals that activate kidney stem cells are not known. We show here that an innate immune, cytokine response after tubule injury is required and sufficient to induce adult zebrafish kidney regeneration. An injury reporter zebrafish transgenic, *Tg(kim1:mScarlet3),* revealed that tubule injury occurred specifically in kidney proximal tubules and was associated with a rapid accumulation of neutrophils and macrophages. Injury also activated a *Tg(NFkB:GFP)* reporter transgene specifically in kidney tubules where RNA seq revealed NFkB target gene and cytokine expression. Inhibition of NFkB signaling with JSH-23 blocked *Tg(NFkB:GFP)* reporter activation and also inhibited induction of new nephrons. Systemic injection of the immune activators lipopolysaccharide or zymosan into uninjured fish rapidly induced cytokine expression followed by nephrogenic gene expression and the appearance of new, functional nephrons. Analysis of injury-induced cytokines revealed that several paralogs of *cxcl11* were strongly expressed throughout the regeneration response and injection of recombinant Cxcl11 was sufficient to induce FGF-dependent kidney stem cell aggregation, but not Wnt-dependent epithelial differentiation. Kidney injury in zebrafish expressing a neutrophil dominant negative *rac2D57N* transgene activated Fgf signaling but failed to induce *wnt9b* or downstream Wnt target genes. Nephrogenic gene expression and epithelial tubule formation was rescued by treatment with the canonical Wnt agonist CHIR. Our findings demonstrate that an injury-induced, sterile immune response regulates kidney regeneration by establishing a nephrogenic niche of Fgf and Wnt signaling that supports tissue-resident kidney stem cell differentiation into functional nephrons.

## Introduction

Injury to mammalian epithelial organs can be repaired by epithelial cell dedifferentiation, proliferation, and redifferentiation to cover denuded basement membranes and regenerate epithelial structure and function ^1,2^. In the zebrafish and other cold-blooded vertebrates, regeneration can also occur from newly forming epithelia ^3,4^, derived from adult progenitor cells ^5–8^. In the zebrafish kidney, injury induces adult kidney stem cells to differentiate into new, functional nephrons ^6^ which engraft and integrate into the existing tubular network ^9^. Zebrafish adult kidney stem cells exist as a scattered mesenchymal cell population in the kidney interstitium, similar in gene expression profile to fetal mouse metanephric mesenchyme ^7,10,11^. and adjacent to kidney distal nephrons. After cessation of kidney growth, these cells are quiescent and function as a reserve cell population that are activated in response to acute injury ^6,10,12^.

Like mammalian kidney tubules, zebrafish kidney cells can be injured by the nephrotoxic antibiotic gentamicin which is endocytosed by proximal tubule cells, accumulates in lysosomes, and triggers lysosome rupture and cell death ^13–16^. Acute tubule injury signals a regenerative response where quiescent stem cells are activated, migrate, and coalesce into rosette-shaped epithelial aggregates on intact distal tubules ^6,12^. Subsequently, kidney stem cell aggregates proliferate, elongate, and differentiate into new epithelial nephrons ^12^. Several families of growth factors and morphogens have been shown to orchestrate regenerative nephrogenesis in adult zebrafish. Fibroblast growth factors *fgf4* and *fgf10a* are induced by acute injury and act as chemokines to recruit kidney stem cells to distal tubules, prior to differentiation into nephron tubules ^17^. Wnt ligands are induced in distal tubules (*wnt9a* and *wnt9b*) and in new stem cell aggregates themselves (*wnt4*) ^12^ and are essential for epithelialization of stem cell aggregates and proliferative nephron outgrowth ^9^. Acute kidney injury also activates renal interstitial cells to secrete prostaglandin E2 (PGE2) which stabilizes β-catenin in kidney stem cell aggregates ^18^. This sequence of events resembles the process of fetal kidney development in mice where Wnt9b expression on the branching ureteric epithelium induces a mesenchymal to epithelial transformation of the metanephric mesenchyme to generate kidney nephrons that subsequently engraft and integrate with the ureteric collecting system ^19^. While the similarities between developmental mammalian nephrogenesis and regenerative nephrogenesis in the zebrafish are becoming clear, it remains unknown how acute kidney injury in zebrafish activates morphogen expression and stimulates kidney stem cells to form new kidney tissue in the context of an adult, functioning organ.

Production of cytokines and other secreted signaling molecules is a well established response to wounding and cell injury in organisms ranging from planarian flatworms to human skin ^20–23^. In zebrafish, organ and tissue damage signals a sterile inflammation, cytokine response that is associated with regenerative responses in the fins, eye, heart, and spinal cord, for example ^24–29^. Evidence for oxidative stress and sterile inflammatory responses after acute zebrafish kidney injury has been reported ^30^ however whether cytokines or immune cell signaling play important roles in zebrafish kidney regeneration remains unknown.

In this study, we investigated signals that initiate regenerative nephrogenesis after kidney tubular cell damage. We find that gentamicin-induced acute kidney injury elicits a robust inflammatory cytokine response which is both required and sufficient to stimulate growth factor expression and initiate adult stem cell-based kidney regeneration. In addition, we find that the cytokine Cxcl11 is rapidly induced in injured tubules and is sufficient to initiate expression of *fgf4* and *fgf10a* and kidney stem cell aggregation on distal tubules, the first stage of new nephron formation. Independently, a neutrophil migration signal induces *wnt9b* expression to drive epithelial differentiation and nephrogenesis. These findings support a model in which inflammatory signaling drives sequential stages of nephron regeneration through a CXCL11–FGF axis and a neutrophil-dependent Wnt signal.

## Results

### Acute kidney injury stimulates inflammatory signaling

*lhx1a:GFP+* zebrafish adult kidney stem cells are activated by acute kidney injury in a step-wise fashion after injury to first aggregate on distal tubules in new nephron rosettes under the control of FGF signaling ^17^, then proliferate under control of Wnt and prostaglandin signaling ^12^, and subsequently differentiate and fuse with existing distal tubules ^9^ to add or replace nephron capacity in the zebrafish kidney (figure 1). In the mouse, Kim-1/Havcr-1 expression marks injured tubule cells ^31^ and is also expressed in macrophages ^32^. To clarify the site of initial tubule damage in gentamicin injured zebrafish we generated a transgenic line incorporating the zebrafish *kim1/havcr1* promoter region (-1,917 kb) driving mScarlet3 expression. *kim1:mScarlet3* expression in kidney epithelia and phagocytes was differentiated by injection and uptake of 10kD fluorescent dextran, with myeloid cell/phagocytes staining double positive for dextran and mScarlet3 (figure 2). Uninjured *Tg(kim1:mScarlet3)* adult kidneys showed low levels mScarlet3 expression in kidney tubules and scattered expression in interstitial, dextran+ cells (figure 2A) representing phagocytes/myeloid cells. Gentamicin injury induced a dramatic increase in kidney tubule *kim1:mScarlet3* expression at 2, 4, and 10 dpi (figure 2 B-D). Consistent with prior reports, acute tubule injury induced an increase in myeloid cells (mScarlet3/dextran double positive) adjacent to injured tubules (arrowheads in figure 2C) ^33^. Gentamicin acute tubule injury was segment-specific and concentrated in the proximal tubule (figure 2D), similar to mammalian kidneys ^13^. Both *Tg(mpeg1.1:GFP)^+^*macrophages and *Tg(mpx:GFP)^+^* neutrophils were recruited to kidney tubules by injury (figure 2 E,F). *Tg(mpx:GFP)^+^*neutrophils showed evidence of fragmentation and in some cases appeared to penetrate epithelial tubules (figure 2F; inset). An increase in kidney myeloid cell number was confirmed by cell counts and whole kidney qRTPCR with myeloid cell marker genes (supplemental figure 1).

**Figure 1.**
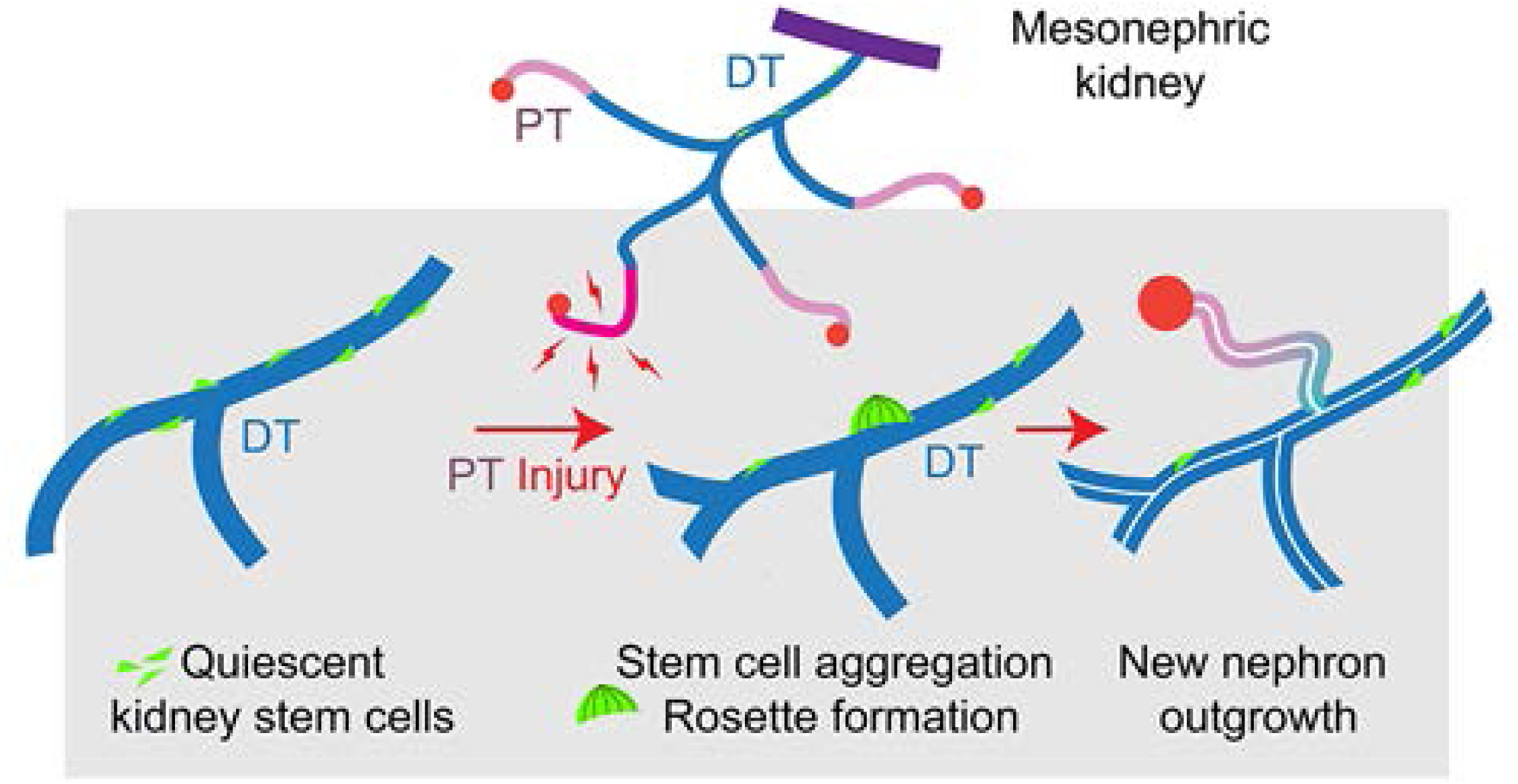
Stages of zebrafish kidney regeneration. The mesonephric kidney is a system of segmented, branched tubules required for blood filtration (glomeruli, red circles), tubular transport (Proximal tubule; PT, Distal tubule; DT), and osmoregulation. Quiescent kidney stem cells (green) reside on distal tubules and in kidney interstitium (left). Injury to the proximal tubule (PT; see figure 2) induces stem cell aggregation on distal tubules under control of fibroblast growth factors, and epithelial nephron outgrowth under control of Wnt signaling ^12,17^.

**Figure 2.**
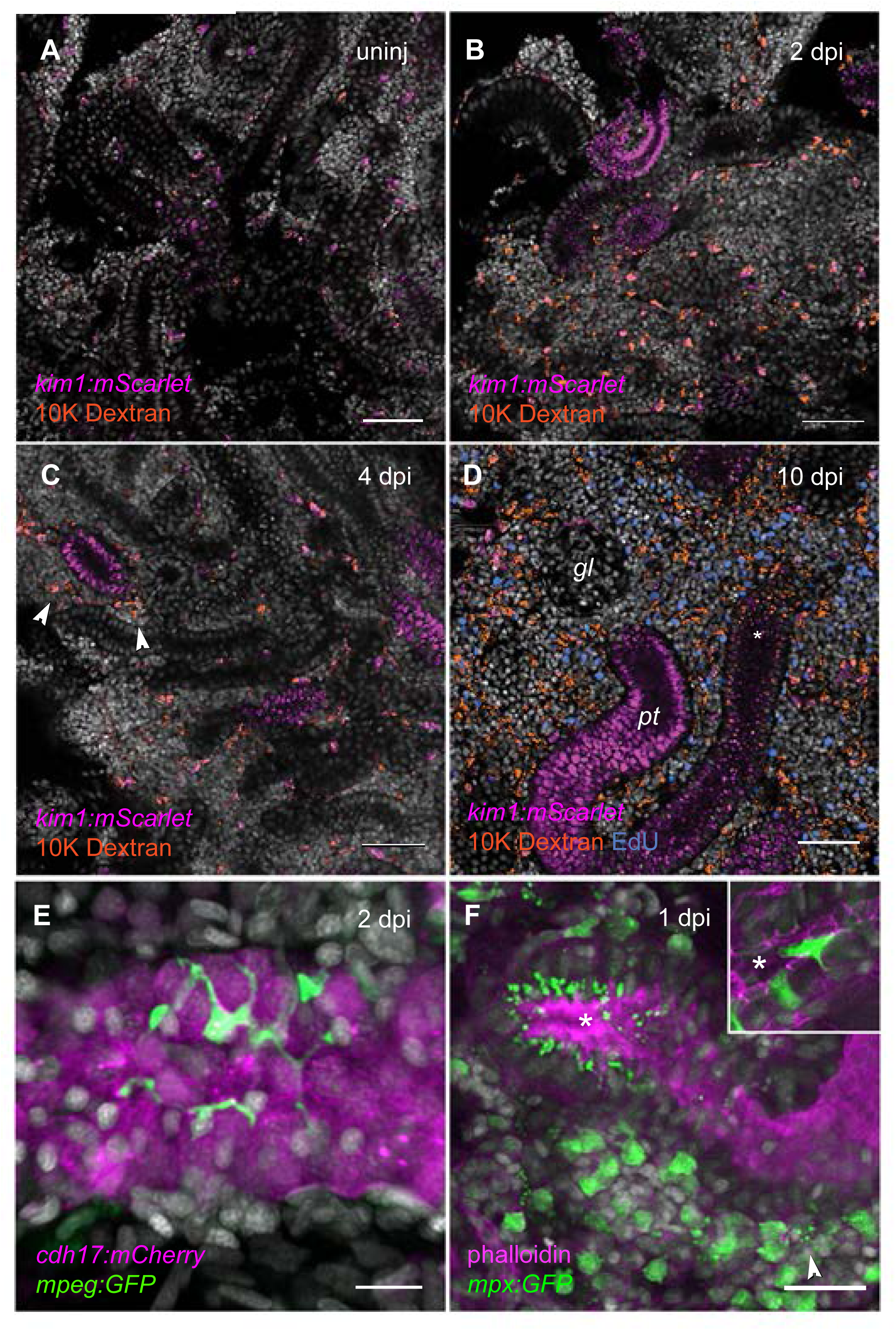
*kim1:mScarlet3* expression marks nephrotoxin-injured tubule segments. (A) Uninjured adult mesonephros with scattered interstitial *kim1:mScarlet3+,* 10K dextran+ phagocytic myeloid cells. (B) At 2 days post injury (dpi), tubule segments express *kim1:mScarlet3* as a marker of tubule damage. (C) At 4 dpi, *kim1:mScarlet3+* tubules are associated with dextran+ phagocytes (arrowheads). (D) A 10 dpi nephron showing the proximal tubule (*pt*) adjacent to a glomerulus (*gl*) is most strongly *kim1:mScarlet3* positive while more distal cells (asterisk) do not express the injury marker. (E) *mpeg:GFP+* macrophages associate with injured *cdh17:mCherry+* tubules at 2 dpi. (F) *mpx:GFP+* neutrophils are ubiquitous in the interstitium and show evidence of fragmentation (arrowhead) in 1 dpi kidneys. Epithelial tubule cells show uptake of GFP. (F, inset) *mpx:GFP+* neutrophils are observed invading epithelial tubule lumens (asterisks) at 1 dpi. Scale bars: A-D = 50 µm; E = 10 µm; F = 20 µm.

To identify signaling pathways associated with injury responses, we performed bulk RNA seq on isolated kidney tubules one and two days after gentamicin nephrotoxin injury (figure 3). We chose bulk RNA seq over the single cell seq approach to generate greater sequence read depth and identify injury-associated rare transcripts in tubules and tubule-associated cells. Fish kidneys are comprised of both nephron tubules and the so-called kidney marrow, the major hematopoietic tissue in the adult fish ^11^. To distinguish RNA transcripts in injured kidney tubule-associated cells from bulk kidney marrow, the tubule fraction was isolated by collagenase digestion, filtering, and sequential washes. Isolated this way, tubule fractions may contain tightly associated immune infiltrating cells (figure 2 E,F) but were significantly depleted of general marrow cells, showing a 50-400-fold depletion of known lymphocyte and hematopoietic marker genes *gata1a, aqp1a, prf1, npsn, hbbe2, and lyz* (figure 3B; supplemental table 1).

**Figure 3.**
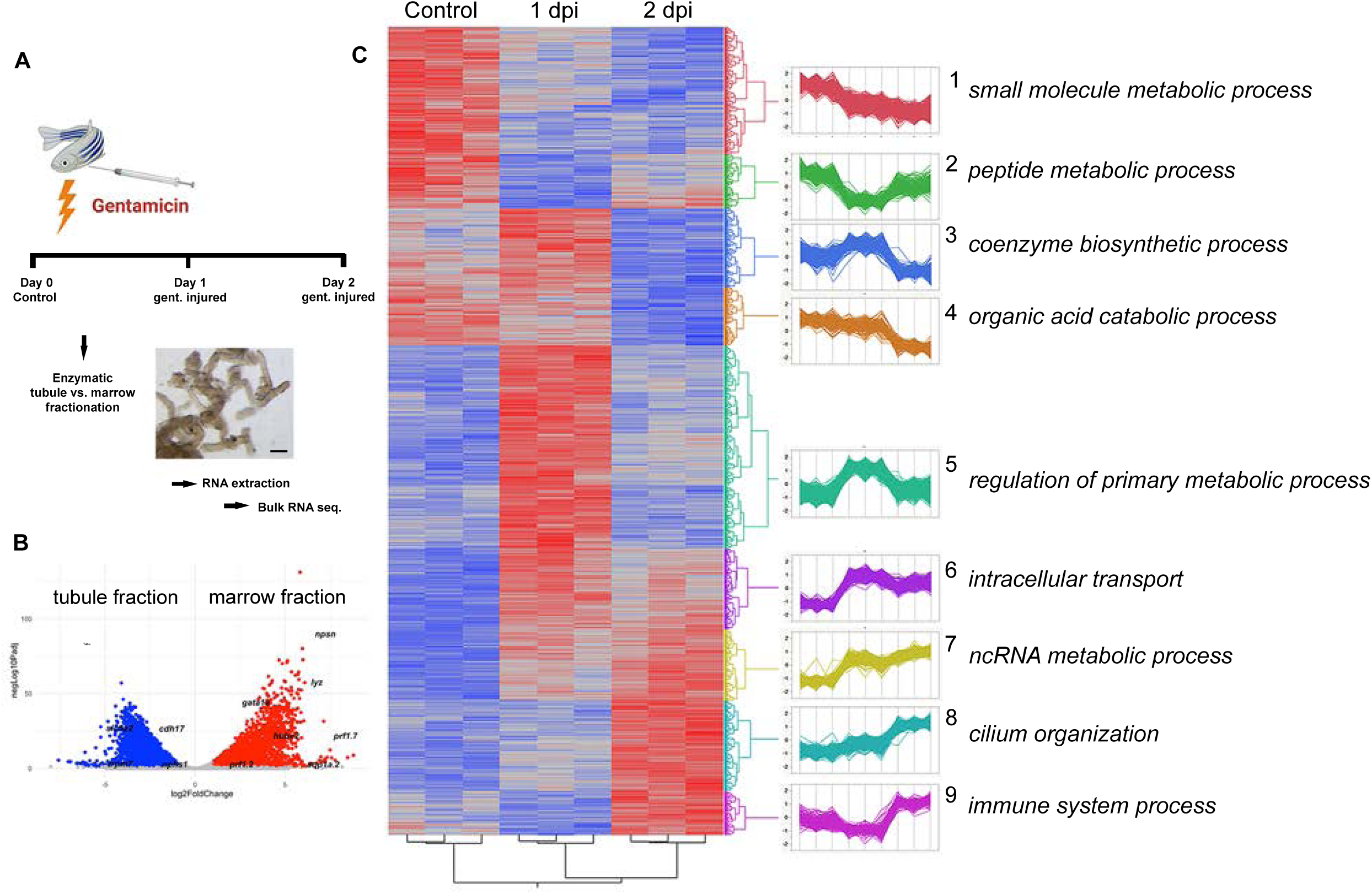
Transcriptomic analysis of acute tubule injury. (A) Injury and sample protocol for tubule purification and sequencing. (B) Volcano plot of tubule (blue) vs. hematopoietic (red) cell fraction transcriptomes. (C) Heat map of kidney tubule transcriptional response to acute injury. Co-regulated transcript clusters and associated GO terms. Inset scale bar = 50 µm.

Analysis of kidney tubule-enriched gene expression at 0-, 1-, and 2-days post injury (dpi) revealed complex expression patterns of multiple co-regulated gene cohorts (figure 3C). As expected from the injury to proximal tubule cells seen figure 2, gene cohorts associated with GO terms related to metabolism, transmembrane transport, and protein synthesis were downregulated on day 1 and day 2 after acute injury (clusters 1, 2 and 4, figure 3, supplemental table 2,3). One day post injury was characterized by a transient upregulation of transcription factors including the immediate-early transcription factor *egr4*, ciliogenic factor *rfx1*, and proinflammatory factor *nfkb2* (supplemental table 2,3). Intracellular and nuclear transport as well as RNA turnover and processing showed sustained upregulation one day post injury (clusters 6,7, supplemental table 2,3). Interestingly, genes associated with ciliogenesis were markedly upregulated at day two (cluster 8, supplemental table 2,3), consistent with *rfx1* induction at day one and prior reports on ciliogenic gene expression after injury in zebrafish larvae ^34^ and post kidney ischemia-reperfusion injury in the mouse ^1^. Notably, inflammatory signaling responses dominated GO terms for genes upregulated in tubule fractions two days post injury (2dpi, cluster 9, supplemental table 2,3). These genes included secreted cytokines (*ccl19, cxcl11.6, ccl34b.8*), interferons (*ifnphi2, ifnphi3*), interferon-induced genes (*ifit8,9,10*), and inducible interferon regulatory factors (*irf10*). Upregulation of the chemotactic factors *cxcl8* (neutrophils) and *cxcl11* (macrophages) was consistent with recruitment of neutrophils and macrophages to injured tubules (figure 2E,F). In addition, HOMER analysis of shared DNA regulatory elements in the genes encoding cluster 9 transcripts revealed enrichment for interferon regulatory factor (*irf*) motifs (supplemental table 4). Early cytokine expression was confirmed by qRTPCR analysis of isolated kidney tubules which revealed strong upregulation of inflammation associated genes two days after gentamicin injury (supplemental figure 2).

The *Tg(cdh17:mCherry)* transgene specifically marks kidney tubule cells in zebrafish larval and adult kidneys ^6^. To confirm inflammation-associated gene expression in tubule epithelial cells, control and 7dpi *Tg(cdh17:mCherry)* FACS tubule cells were analyzed for genes upregulated by injury. Consistent with day 1 and day 2 post injury results, day 7 injured FACS tubule cells showed strong upregulation of inflammation and interferon regulated genes and cytokines associated with top gene ontology terms immune response (GO:0006955) and defense response (GO:0006952) (supplemental table 5). Also consistent with day 1 and 2 results, genes upregulated by injury included multiple *cxcl11* paralogs (*cxcl11.1, cxcl11.5, cxcl11.6, cxcl11.8).* The Cxcl11 receptors, *cxcr3.1, cxcr3.2, cxcr3.3* ^35^, were also expressed in *Tg(cdh17:mCherry)+* tubule cells (supplemental table 5). Overall, RNA seq revealed that damage-associated sterile inflammation is a persistent and conserved feature of acutely injured zebrafish kidney epithelial tubules and tubule-associated cells.

### Inflammatory signaling is required and sufficient for kidney regeneration

The zebrafish *Tg(NF-kB:GFP)* reporter line contains a synthetic enhancer element consisting of 6 repeats of an NFkB binding site and reports in vivo transcriptional activity of NFkB ^36^. Upregulation of cytokines observed kidney tubule fractions after gentamicin injury correlated with induction of *Tg(NF-kB:GFP)* reporter expression in injured tubules (figure 4A,B). *Tg(NF-kB:GFP)* reporter activity was highest in proximal tubules adjacent to glomeruli (figure 4B) and was excluded from *Tg(slc12a3:mCherry)+* distal tubules ^37^ (supplemental figure 3). These results are consistent with injury induction of *Tg(kim1:mScarlet3)* expression specifically in proximal tubules (figure 2) and imply that injury signaling must act at a distance to initiate new nephron formation on distal tubules.

**Figure 4.**
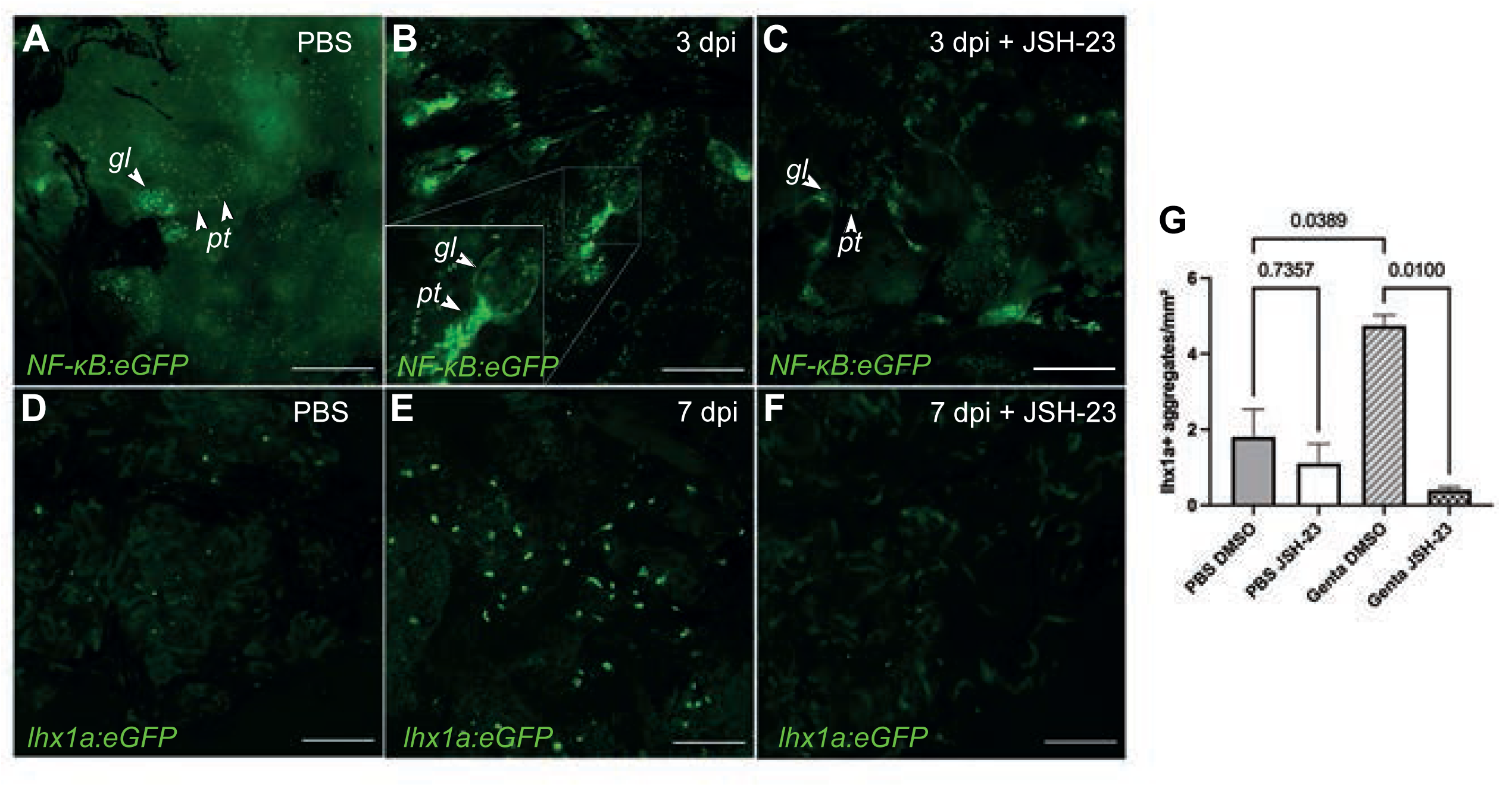
NF-kB signaling is required for kidney regeneration. (A) Uninjured *NF-kB:GFP* reporter transgenic kidney tissue. gl: glomerulus, pt: proximal tubule. (B) Upregulation of *NF-kB:GFP* reporter activity in proximal tubule segments (pt). (C) Incubation with NF-kB nuclear translocation inhibitor JSH-23 prevents activation of *NF-kB:GFP* reporter 3 days post injury. (D) Uninjured *lhx1a:eGFP* (kidney stem cells) transgenic kidney tissue. (E) *lhx1a:eGFP+* new nephron cell aggregates 7 days post gentamicin injury. (F) *lhx1a:eGFP* transgenic kidney tissue 7 days post gentamicin injury treated with JSH-23. (G) Quantification of effect of JSH-23 on kidney regeneration. Scale bars = 200 µm.

To determine whether NFkB activity was required for stimulating kidney progenitor cells, we blocked NF-kB nuclear translocation with the small molecule drug JSH-23 ^38^ during the response to acute injury. Treatment of gentamicin injured *Tg(NFkB:GFP)* fish with JSH-23 prevented activation of the *NFkB:GFP* reporter at 4 dpi (figure 4C), confirming its pharmaceutical efficacy. Notably, JSH-23 also prevented the appearance of *Tg(lhx1a:GFP)^+^* nephron progenitor cell aggregates at 7 dpi (figure 4D-G). The results indicate that NFkB transcriptional activity is required for *lhx1a*+ nephron cell aggregate formation in zebrafish kidney regeneration.

To test whether inflammation and cytokine signaling may be sufficient to induce kidney regeneration in the absence of tubule injury, we injected adult zebrafish intraperitoneally with the pathogen-associated molecular pattern inflammatory stimuli Lipopolysaccharide (LPS), zymosan, and polyI:C and assayed cytokine and nephrogenic gene expression. As expected, LPS, Zymosan, or polyI:C injection induced a rapid and transient expression of cytokines *il6, il-1ß, tnf-a* and the inflammatory response genes *mxa* and *pim-1* in whole kidney that peaked at three to six hours post injection and returned to baseline by 48 hours post injection (supplemental figure 4). Compared with the gentamicin inflammatory response, LPS induction of cytokines was rapid and transient while gentamicin induced a slower, more prolonged response to at least 4 dpi (supplemental figure 4). Also, LPS strongly induced *il6, il-1ß, and tnf-a* expression that was not significantly observed in gentamicin injected fish (supplemental figure 4), implying different inflammatory pathways are activated in these two conditions. Notably, LPS injection significantly induced NFkB reporter activity and the nephrogenesis markers *lhx1a*, and *fzd9b* at seven days post injection, without significantly altering the expression of injury markers *kim1/havcr1* or *slc20a1a* (figure 5A). Also, LPS did not induce expression of the *kim1:mScarlet3* injury reporter (supplemental figure 5) indicating that induction of nephrogenesis markers was unlikely to be a response to an LPS-induced tubule injury. Expression of *wnt9b* mRNA, a distal tubule morphogen required for nephrogenic regeneration ^9,12^, was also induced by LPS injection (figure 5A). In addition, zymosan, but not polyI:C, induced significant expression of *lhx1a* at 7 days post injection (supplemental figure 6), consistent with reports that zymosan and polyI:C engage different Toll-like receptors and signal distinct immune responses ^39^.

**Figure 5.**
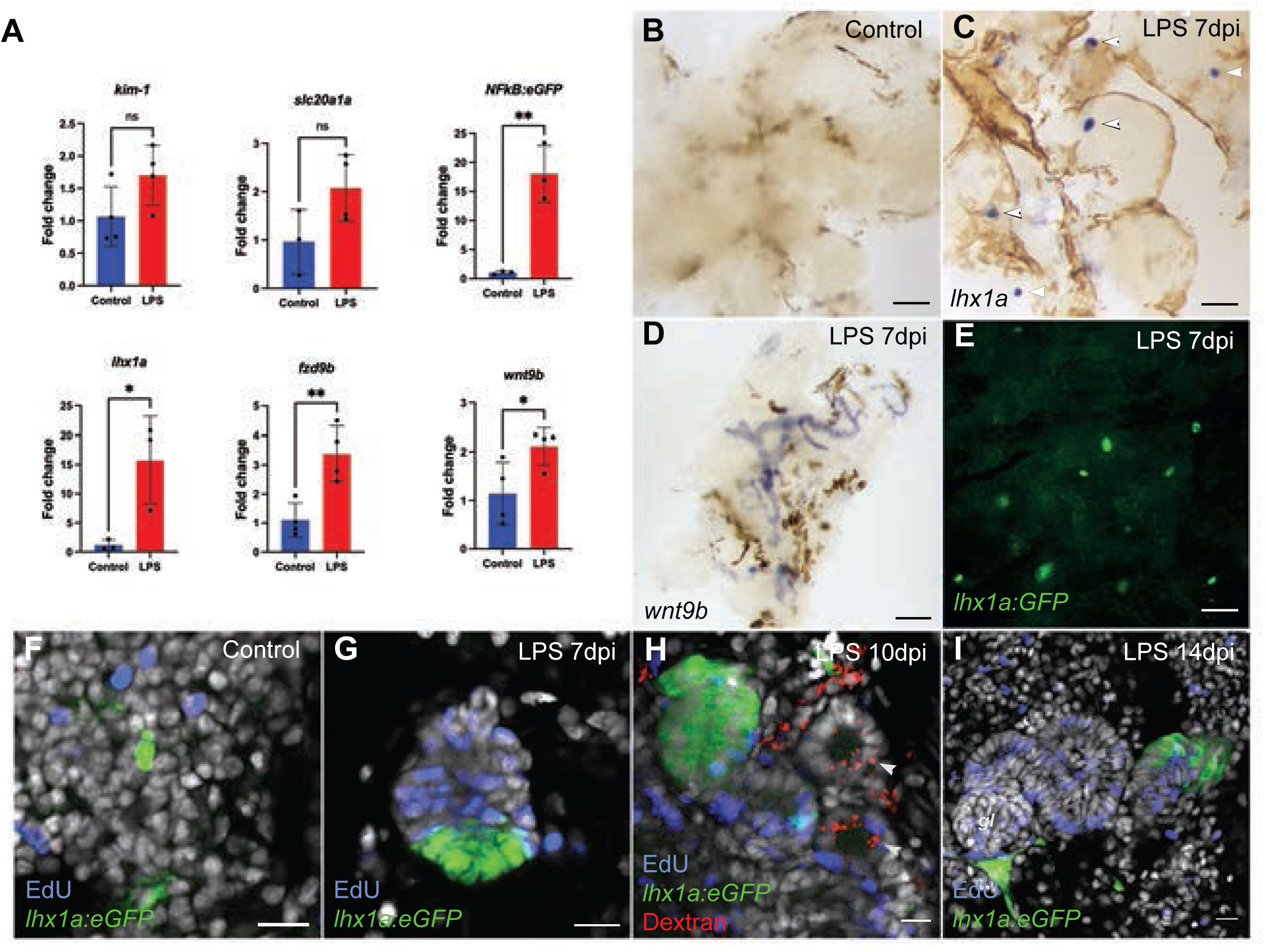
Induction of nephrogenesis by LPS without injury. (A) Quantification (qRTPCR) of NFkB reporter and nephron marker gene expression, seven days post LPS injection. (B) Control PBS injection kidney shows no nephrogenesis. (C) Seven days post LPS injection, new *lhx1a+* nephron stem cell aggregates have formed (arrowheads; in situ hybridization). (D) Seven days post LPS injection, kidney distal tubules and collecting ducts express *wnt9b.* (E) Transgenic lhx1a:eGFP kidneys show lhx1a promoter reporter activity in new nephrons seven days post LPS injection. (F-I) Time series of nephrogenesis after LPS injection. (F) Uninjected fish show single, non-proliferating *lhx1a:eGFP+* kidney stem cells. (G) Seven days post LPS injection, *lhx1a:eGFP+* proliferating (EdU: blue) new nephron cell aggregates have formed. (H) By 10 days post LPS, labelled dextran (red) filtration can be seen in new nephrons marked by EdU incorporation (blue). (I) By 14 days post LPS, fully formed, Edu marked new nephrons show persistent distal *lhx1a:eGFP*+ expression and morphologically complete glomeruli (*gl*) at the proximal nephron end. Scale bars: B-E = 200 µm; F,G = 20 µm; H,I = 10 µm.

Wholemount in situ hybridization demonstrated that LPS-induced expression of *lhx1a* occurred in new nephron cell aggregates (figure 5 B,C). LPS also induced *wnt9b* expression specifically in adult kidney tubules (figure 5D) at 7 dpi. LPS-induced *Tg(lhx1a:GFP)^+^* nephrons (figure 5E) were indistinguishable from nephrons induced by acute kidney injury (figure 4; ^9^). At higher resolution, uninjected control kidneys showed scattered EdU+ interstitial cells and single, non-proliferating *lhx1a:eGFP+* kidney progenitor cells (figure 5F). By 7 days post LPS injection, proliferating, EdU+, *lhx1a:GFP*+ aggregates were observed (figure 5G). Intraperitoneal injection of 10KD fluorescent dextran demonstrated that LPS-induced, EdU+ new nephrons took up dextran from epithelial tubule lumens (figure 5H) at 10 dpi, an indicator of functional glomerular fluid filtration. LPS induced nephrons at 14 dpi had fully formed glomeruli and remained highly proliferative (figure 5I). The results indicate that inflammatory signaling may be sufficient by itself to induce fully functional nephrons without causing significant tubule injury.

### Kidney regeneration is initiated by a Cxcl11 - Fgf signaling cascade

Both gentamicin injury and LPS injection induced nephrogenesis, suggesting either that LPS may be inducing a tubule injury below the level of detection by *kim1* or *slc20a1a* expression changes, or that cytokines generated by both treatments may be the diffusible signals that initiate kidney regeneration. To test this we performed initial screen of recombinant Tnfa, Il1-b, Il-6, or Clcf1 peptide injections, however none of these cytokines induced nephrogenic marker gene expression (supplemental figure 7). To better define cytokine candidates that may induce nephrogenesis, we examined the intersection of LPS ^40^ and our gentamicin-induced cytokine expression profiles. The zebrafish cytokines *ccl34a, ccl35, tnfa, cxcl11, and cxcl18b* were found to be induced in common between LPS and gentamicin injury and therefore candidates for inducers of nephrogenesis. Since Tnfa was negative in our prior experiments (supplemental figure 7), we pursued other candidate cytokines. Cxcl11 is 1) known to act as a chemokine for macrophages ^35^, 2) is a transcriptional target of NFkB ^41^, and multiple zebrafish *cxcl11* paralogs were upregulated by injury in tubule and tubule-associated cells in response to tissue damage (this study, supplemental figure 2, supplemental table 6). Since *cxcl11* and related cytokines can also be expressed by myeloid cells that amplify initial cytokine responses ^42,43^, we examined *cxcl11.6* expression by in situ hybridization. *cxcl11.6* expression was not detected in control, uninjured kidneys (figure 6A) but was strongly expressed at 4 dpi in patches of interstitial cells (figure 6B) adjacent to kidney epithelial tubules (figure 6C), suggesting that cytokine expression is amplified by non-tubule, immune cells as injury and regeneration responses proceed. Similar results were observed for the paralog *cxcl11.1* (suppplemental figure 8).

**Figure 6.**
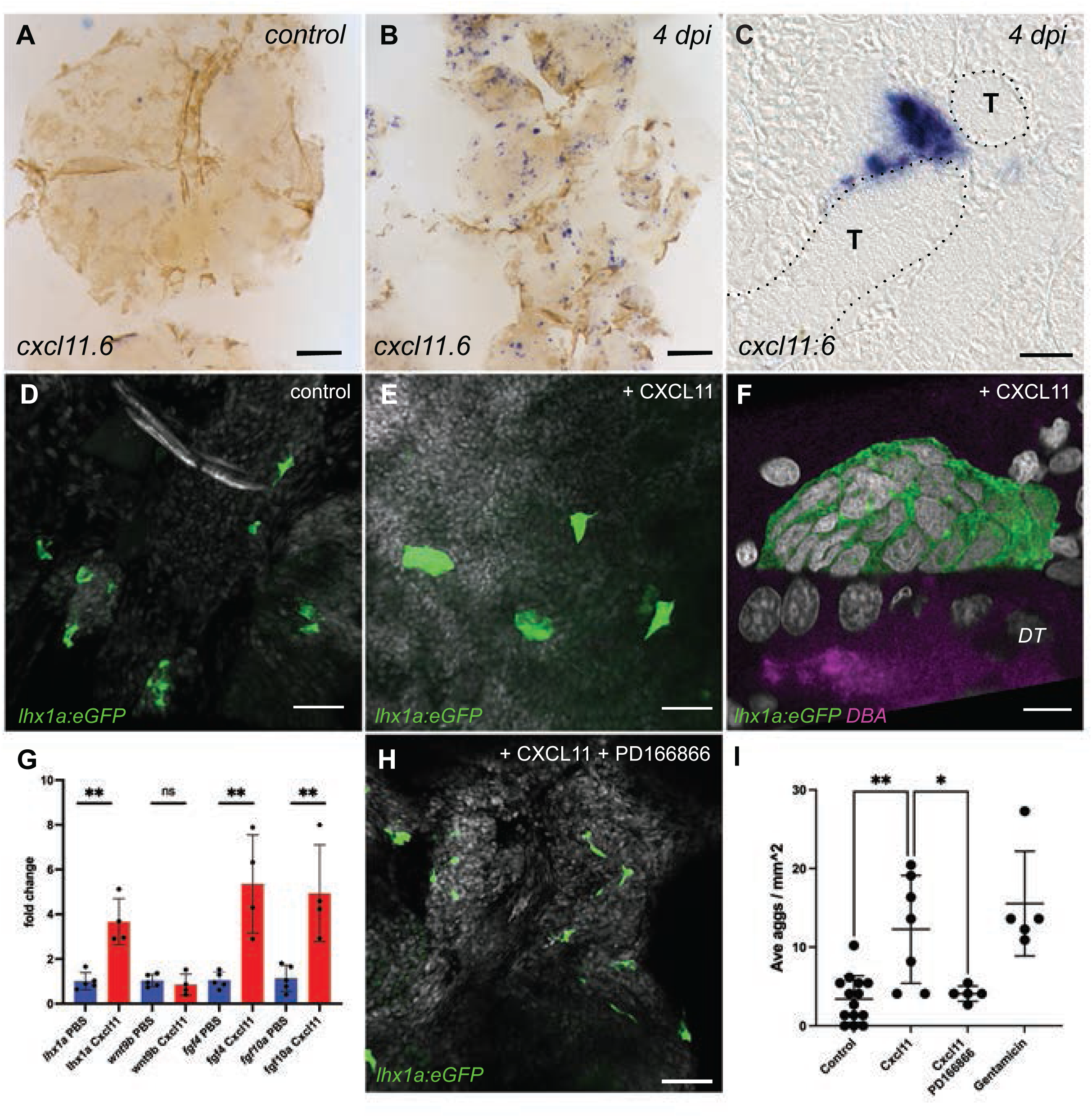
Cxcl11 and FGF expression drive aggregation of kidney stem cells. Whole mount in situ hybridization of control (A) and 4 days post injury (4 dpi) kidneys (B) shows expression of *cxcl11.1* in patches of interstitial cells (C) adjacent to kidney tubules (T in C). (D) Control kidney *lhx1a:eGFP+* single kidney stem cells prior to CXCL11 cytokine injection. (E) three days post CXCL11 injection, multicellular *lhx1a:eGFP+* aggregates form. (F) High magnification view of a multicellular *lhx1a:eGFP+* cell aggregate formed on a dolichus biflorus lectin-lumen (DBA) marked distal tubule (magenta; *DT*). (G) Gene expression changes after CXCL11 injection quantified by qRTPCR. (H) Inhibition of Fgfr1 signaling by PD166866 blocks CXCL11-induced multicellular *lhx1a:eGFP+* cell aggregate formation. (I) Quantification of *lhx1a:eGFP+* cell aggregates formed three days after CXCL11 injection and the effect of Fgfr1 inhibition. Scale bars in A,B = 200 µm; C = 20 µm; D,E,H = 40 µm; F = 5µm.

Prior work has shown that human recombinant CXCL11 is active in zebrafish ^44^, so we tested whether injection of recombinant CXCL11 could induce nephrogenesis in uninjured fish. Remarkably, intraperitoneal injection of recombinant CXCL11 alone was sufficient to induce *lhx1a:GFP+* stem cell aggregation on uninjured adult zebrafish kidney distal tubules, the normal site of new nephron formation ^12^. Single *lhx1a:eGFP+* cells present in control kidneys (figure 6D) were induced to form multicellular aggregates after CXCL11 injection (figure 6E,F,I). These aggregates did not progress to the nephron tubule outgrowth stage however (figure 1), and instead persisted as multicellular domes with no evidence of epithelial cell differentiation or interconnection with distal tubules ^9^ (figure 6E). High resolution confocal imaging confirmed the multicellular nature of the aggregates and showed that they form on dolicus biflorus lectin-positive distal tubules (figure 6F), the normal site of new nephron formation ^12^. The results suggest that Cxcl11 secretion in response to injury is a key initial step in new nephron formation.

Zebrafish kidney stem cell migration and aggregation is mediated by injury-induced expression of *fgf4* and *fgf10a* in kidney tubules and Fgfr1 signaling ^17^. Subsequent cell polarization, epithelial tubule outgrowth, and connection to distal tubules is Wnt-dependent (figure 1) ^9,12^. Significantly, CXCL11 injection induced expression of *fgf4* and *fgf10a* but not *wnt9b* in whole kidney tissue (figure 6G). We tested whether CXCL11-induced stem cell aggregate formation was Fgf-dependent by treating CXCL11-injected fish with the Fgfr1 receptor antagonist PD166866 ^17^. Inhibition of Fgfr1 signaling blocked CXCL11-induced kidney stem cell aggregation (Figure 6H,I), demonstrating that CXCL11 effects are mediated sequentially by induction of fibroblast growth factors. The results provide evidence for a regenerative cell signaling cascade where injury-induced *cxcl11* expression in tubules and/or tubule-associated cells induces *fgf4/10a* expression to signal kidney stem cell aggregation on distal tubules and initiate kidney regeneration.

### Neutrophil motility is required for Wnt signaling in kidney regeneration

Following acute injury, neutrophils are the first cells to arrive at damaged tissue and proceed to recruit macrophages and other immune cells to generate an injury/wound healing response ^45^. We noted a significant increase in myeloid cell number in association with tubule injury (figure 2, supplemental figure 1), similar to what has been reported previously in zebrafish injury models ^33^. We tested whether impaired neutrophil migration in the neutrophil-specific dominant negative Rac2 line *Tg(mpx:mCherry,rac2_D57N)* ^46^ would impact cytokine production or subsequent regenerative responses in kidney injury. *Tg(mpx:mCherry,rac2_D57N)* expressing kidneys showed normal *cxcl11*, *fgf4,* and *fgf10a* induction after injury (figure 7A-C). *dusp6,* an FGF target gene was also normally induced after injury (figure 7D), indicating that neutrophil motility was not required for initial *cxcl11-fgf* signaling. However, *Tg(mpx:mCherry,rac2_D57N)* kidneys failed to induce the Wnt ligand *wnt9b* and the Wnt target genes *lhx1a, lef1 and wnt4* after kidney injury (figure 7E-H), suggesting that neutrophil motility and signaling was required for Wnt-dependent differentiation of new nephrons. Wholemount in situ hybridization (figure 7I-K; supplemental figure 9) and quantification of new nephrons (figure 7L) confirmed that stem cell aggregate formation marked by the Fgf target gene *dusp6* ^17^ occurred normally in injured *Tg(mpx:mCherry,rac2_D57N)* kidneys while whole mount in situ expression of *lhx1a* expression was markedly reduced (figure 7M-O; 7L).

**Figure 7.**
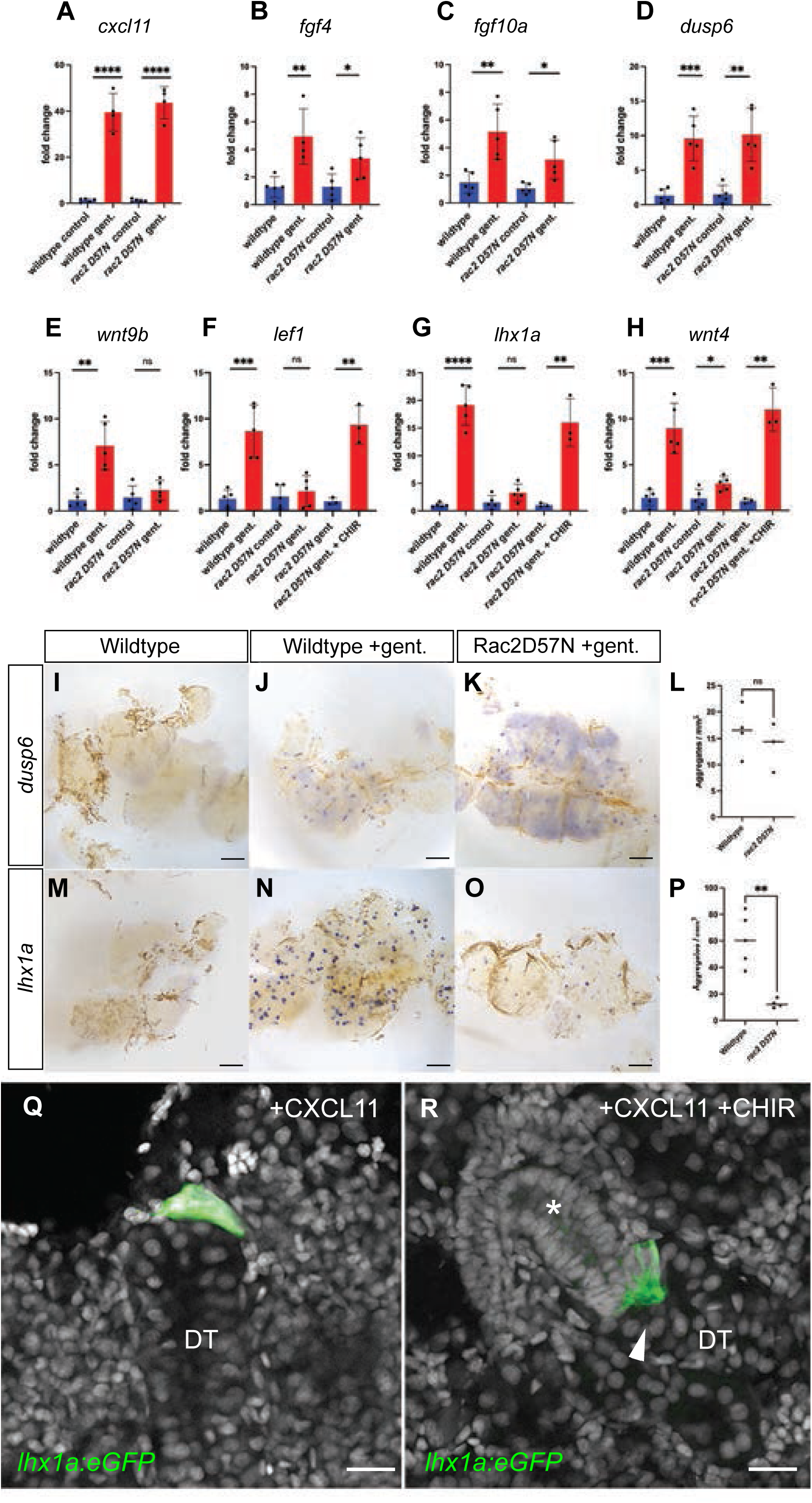
A neutrophil motility-dependent signal is required for Wnt signaling and nephrogenesis. Quantification of 2 dpi *cxcl11.6* (A), 3 dpi *fgf4* (B), 3 dpi *fgf10a* (C), and 7 dpi *dusp6* (D) signaling by RTPCR shows no significant difference from controls in the *Tg(mpx:mCherry,rac2_D57N)* transgenic after kidney injury. Quantification of 7 dpi *wnt9b* (E), 7dpi *lef1* (F), 7dpi *lhx1a* (G), and 7 dpi *wnt4* (H) expression after injury shows a lack of *wnt9b* induction by injury and a lack of response of Wnt target genes in the *Tg(mpx:mCherry,rac2_D57N)* transgenic that can be restored by treatment with 10µM CHIR from 4-6 dpi. (I-L) Whole mount in situ hybridization for *dusp6*. Compared to control uninjured kidney (I), gentamicin acute injury induces multiple new nephron cell aggregates that are unaffected in the *Tg(mpx:mCherry,rac2_D57N)* transgenic (K, L). (M-P) Whole mount in situ hybridization for *lhx1a*. Compared to control kidney (M), gentamicin acute injury (N) induces many *lhx1a+* new nephron aggregates. In cell aggregates of *Tg(mpx:mCherry,rac2_D57N)* transgenic kidneys (O), *lhx1a* is not highly expressed. Quantification of aggregates in N and O shows a significant reduction in *lhx1a+* new nephron aggregates in the *Tg(mpx:mCherry,rac2_D57N)* transgenic. (Q) Injection of CXCL11 in wildtype zebrafish generated new nephron aggregates marked by *lhx1a:eGFP* that did not elongate into nephron tubules. (R) Treatment of CXCL11 injected zebrafish with 10 µg/ml CHIR between 4 and 6 days induced tubule outgrowth (asterisk) and distal tubule invasion (arrowhead) in a subset of treated animals. Scale bars: I-O = 200 µm; Q,R = 20 µm.

Both gentamicin-injured *Tg(mpx:mCherry,rac2_D57N)* and CXCL11-injected fish generated nephron stem cell aggregates that did not progress to the Wnt-dependent epithelial differentiation phase of kidney regeneration. We therefore tested whether supplementing acutely injured *Tg(mpx:mCherry,rac2_D57N)* or CXCL11-injected kidneys with the canonical Wnt agonist CHIR ^47^ would restore nephrogenic Wnt target gene expression and promote tubulogenesis. *wnt9b* induction after acute injury normally occurs in distal tubules and peaks at five days post injury ^12^. *Tg(mpx:mCherry,rac2_D57N)* gentamicin-injured or CXCL11-injected fish were treated with CHIR for 48 hours between 4 and 6 dpi and assayed for nephrogenesis at 7 dpi. Expression of Wnt target genes *lhx1a, lef1,* and *wnt4* was completely restored by CHIR treatment in *Tg(mpx:mCherry,rac2_D57N)* gentamicin-injured kidneys as assayed by qRTPCR (figure 7F-H). CHIR treatment also induced epithelial cell differentiation and tubule outgrowth from *lhx1a:eGFP+* new nephron aggregates in CXCL11-injected fish, imaged by confocal microscopy (figure 7Q,R). Quantification of nephron tubule formation frequency revealed that while aggregates induced by CXCL11 injection alone never formed nephron tubules, 16.7% (7/42) of CXCL11 +CHIR treated *lhx1a:eGFP+* aggregates showed evidence of epithelial differentiation and tubule outgrowth (figure 7R). Elongating nephrons in CXCL11 +CHIR treated fish also showed evidence of distal tubule invasion, a Wnt-dependent process ^9^, as a prelude to tubule interconnection and functional engraftment (figure 7R).

## Discussion

An important aim in regenerative biology is to determine how organ repair can be achieved without fibrosis and prevent chronic deterioration of organ function. Regenerative responses to injury in epithelial organs are markedly different between fish, that exhibit scarless healing, and mammals that develop fibrotic, degenerative responses to repeated organ injury ^48–50^, making fish and other cold-blooded vertebrates important systems for understanding organ regeneration ^51,52^. While it is increasingly clear that inflammatory responses and cytokine secretion are universal responses to wounding and injury ^53^, whether inflammation induces beneficial or damaging responses in injured tissue and how cytokine signaling may coordinate responses of tissue progenitor cells to stimulate regeneration is not well defined.

The generation of wound and injury signals as an initial stimulus for regeneration is well conserved in species ranging from humans to planaria ^20,54–56^. In zebrafish, inflammatory signaling has been identified as a positive regulator of regeneration in multiple tissues, including the fin, retina, brain, spinal cord, and heart ^57–62^. In the zebrafish kidney, injury stimulates the production of new nephrons from tissue-resident stem cells to compensate for lost organ function ^3,6–8,12,17^. We show here that inflammatory signaling is essential for this response. We also find that stimulating inflammation in the absence of tissue injury is sufficient to generate fully functional nephrons. Our results show that full regeneration requires multiple, independently generated inflammatory signals acting in combination to stimulate nephrogenesis: injury induction of *cxcl11* leads to expression of fibroblast growth factors *fgf4* and *fgf10a* which is sufficient to recruit kidney stem cells to the distal tubules and form pretubular cell aggregates ^17^. An independent, neutrophil-associated inflammatory signal is essential for subsequent *wnt9b* expression, epithelial tubule differentiation ^12^, and engraftment to the distal tubules ^9^. This inflammation-induced sequence of Fgf and Wnt growth factor signaling parallels developmental signaling in mammalian nephrogenesis ^63,64^ and the protocols used for human kidney organoid production ^65,66^, indicating that both developmental and inflammatory regulatory circuits can stimulate nephron formation by generating a nephrogenic niche enabled by induction of critical growth factors.

Inflammatory signaling in the context of injury is typically biphasic: damage signals drive macrophage and neutrophil recruitment to sites of injury, supporting apoptotic cell clearance, and stimulating regenerative responses that are all subsequently downregulated as part of productive repair ^67^. Persistent, unresolved inflammation drives fibrotic, degenerative tissue responses that result in failed repair and pro-inflammatory senescent cell states ^1,68^. The difference between these outcomes may be more dependent on the magnitude and duration of the responses than the identity of signaling molecules themselves ^67,69^. In mammals, Wnt ligands released from macrophages can promote pro-regenerative proliferation of intestinal, hepatic, and renal epithelial cells ^67^, while prolonged exposure of injured tissue to Wnt signals can result in myofibroblast differentiation and fibrosis ^69–71^. In the zebrafish, injury to the heart or the kidney does not result in fibrotic scarring despite the expression of similar, injury-induced ligands ^12,72,73^. Also, we have previously shown that the Wnt ligand *wnt9b* is only transiently upregulated by acute kidney injury, peaking at 5 dpi and resolving at 7 dpi ^12^, suggesting that signaling responses to injury are temporally well controlled in the fish ^74^.

Cellular context is also a determinant of regenerative vs. degenerative outcomes. The presence of different populations of immune cells or tissue-resident organ progenitors is likely to have a major effect on regenerative outcomes. In neonatal mouse hearts that show scar-free healing, an embryo-derived population of resident macrophages promotes cardiomyocyte proliferation and angiogenesis after injury while the adult heart is repopulated post-injury by pro-inflammatory macrophages ^75^. In axolotl limb regeneration, macrophages show different responses to DAMPs compared to the mouse ^76^, suggesting that the interpretation of inflammatory signals is different in regenerative vs. non-regenerative animals. It is not known whether immune cell responses in the zebrafish kidney differ in important ways from the mouse however, the presence of adult kidney stem cells in the fish ^6,7,11^ clearly distinguishes the regenerative potential of the fish kidney from the mouse. It is clear that the same ligand (Wnt9b) can induce divergent responses in the fish kidney (epithelialization of stem cells) compared to the mouse (myofibroblast differentiation), due to different developmental potentials of resident interstitial cells. Sorting out how fish maintain multipotent progenitor cells in the adult kidney ^7^ and how cell lineage potential is determined in fish vs. mouse will be important questions for future studies.

Inflammatory signaling cytokines we describe could act directly on stem/progenitor cells or, alternatively, could act indirectly, creating a secondary signaling environment that recapitulates nephrogenic developmental cues. In the mouse lung, IL-6/STAT3 signaling stimulates regeneration of airway ciliated cells by acting directly on basal stem cells, promoting their differentiation ^77^. Also, in mammalian intestinal regeneration, Il-22 from lymphoid cells can act directly Lrg5+ progenitors to support organoid formation in vitro and promote intestinal tissue regeneration in vivo ^78,79^. On the other hand, Wnt expression from gut macrophages has been shown to generate a pro-regenerative niche that mitigates intestinal epithelial injury ^80^. Evidence in our study points to an indirect mechanism of action of inflammatory cytokines on kidney regenerative responses. The activity of Cxcl11 signaling kidney stem cell aggregation requires Fgfr1 signaling and correlates with induction of the Fgf ligands *fgf4* and *fgf10a* which are normally expressed in distal tubule epithelial cells after injury ^17^. This primary site of new nephron formation ^12^ is distinct from the stem cells themselves. Similarly, the neutrophil motility requirement for epithelial differentiation correlates with expression of *wnt9b,* which normally occurs in distal tubules, not the kidney stem cells. The simplest interpretation of our results is that injury-induced inflammatory signaling acts on pre-existing distal tubules to generate secondary signals (*wnt9b, fgf4/10a*) and create a nephrogenic signaling niche. Consistent with this idea, following injury, kidney stem cells do not proliferate until after they aggregate and begin forming an epithelial tubule at 5-7 dpi (unpublished results and ^7^), implying that cytokines do not act directly as mitogens on stem cells. Our transcriptomic data (supplemental table 5) and data from other studies ^37^ show that the Cxcl11 receptors Cxcr3.1, Cxcr3.2, and Cxcr3.3 are expressed in zebrafish distal tubule cells. However, myeloid cells/macrophages are also known to express Cxcr3, respond to Cxcl11 ^35^, and could act as signaling intermediates promoting kidney stem cell / distal tubule interactions. Further studies of the cell types involved in responses to the inflammatory signals we have identified will better define the sources, direct targets, and signal transduction pathways of cytokines responsible for establishing a nephrogenic niche after acute kidney injury.

## Materials and Methods

### Zebrafish maintenance

Adult wild-type TuAB zebrafish (*Danio rerio*), both male and female, aged approximately 6–18 months, were used for this study. The animals were maintained according to established protocols ^81^. All animal experiments were performed in compliance with MDIBL IACUC animal welfare regulations. Transgenic fish used were: *Tg(lhx1a:eGFP)* ^6^*, Tg(NF-kB:GFP)* ^36^*, Tg(Kim1:mScarlet3)* (see below)*, Tg(mpeg1:eGFP)* ^82^*, Tg(mpx:GFP*) ^83^, and *Tg(mpx:mCherry,rac2_D57N)* ^46^. Experimentally induced acute kidney injury procedures were carried out in accordance with MDIBL-approved IACUC protocols.

### Generation of *kim1:mScarlet3* transgenic

The *kim1:mScarlet3* transgenic line was generated by InVivo Biosystems (Eugene, OR). One cell stage wildtype (ABC) zebrafish embryos were injected with 2 ul of a mixture of the Tol2 *Tg(kim1(-1,917):mScarlet3* donor plasmid (25 ng/μL), Tol2 transposase mRNA (25 ng/μL), and phenol red injection dye (0.05%). F0 injected embryos with mosaic expression of the fluorescent marker consistent with published *havcr1/kim1* in situ hybridization (see https://zfin.org/ZDB-GENE-040718-131/expression) were raised to adulthood and outcrossed to identify transgenic founders. Germline transmission was confirmed by visualization of mScarlet3 expression in putative macrophages in the F2 generation.

### Drug treatments

Fgfr1 signaling was inhibited by treating adult zebrafish with PD166866 (10 μM; prepared from a 25.2mM stock, final DMSO concentration 0.04%) added to system water, as described by ^17^. Fish were pre-treated for 24 hours prior to recombinant CXC11 injection and maintained in the treatment solution for an additional 3 days. Control animals were kept in system water containing an equivalent concentration of DMSO. Both treatment and control solutions were changed daily by replacing half of the volume with fresh drug-treated water. The FGF inhibition protocol was continued until the animals were sacrificed 3 days post-CXCL11 injection. NF-kB signaling was inhibited by treating adult zebrafish with JSH-23 in 500 ml fish system water containing 3.5 μM JSH-23 (final DMSO concentration 0.1% DMSO, Sigma Aldrich J4455) as described in PMID 31866203. Fish treatments started 24 hours post-injection with gentamicin and were carried out for 7 days. Control animals were kept in system water containing an equivalent concentration of DMSO. Both treatment and control solutions were changed daily by replacing half of the volume with fresh drug-treated water. For CHIR treatment, gentamicin injected Rac2DN fish (*Tg(mpx:mCherry,rac2_D57N)* were incubated in 10µg/ml CHIR (CHIR 99021, Tocris cat. #4423) in 0.1% DMSO for 48 hours between 4 and 6 days after injury, after which the medium was replaced with fresh drug-free water and the fish analyzed at 7dpi.

### Intraperitoneal injection

Fish were anesthetized in 0.024% tricaine. Acute kidney injury was induced by intraperitoneal injection of gentamicin (Sigma, Saint Louis, MO) at 80mg/kg (2mg/ml stock in PBS at 10μl per 0.25g body mass) as previously described ^84^. Sham-injured controls received an equivalent volume of PBS in place of gentamicin. Kidney injury was confirmed at 1-day post-injection by visual detection of cast excretion, as previously reported ^84^. For immune stimulation, adult fish were intraperitoneally injected with Lipopolysaccharide (LPS) from *E. coli,* (Sigma, Saint Louis, MO. 10mg/ml stock in PBS at 10μl per 0.5g of body mass), Zymosan from *S. cerevisiae,* (Sigma, Saint Louis, MO. 10mg/ml stock in PBS at 20μl per 0.5g of body mass). Polyinosinic–polycytidylic (poly I:C) (acid Sigma, Saint Louis, MO. 10mg/ml stock in PBS at 25μl per 0.5g of body mass). Fish injected with an equivalent volume of PBS served as controls. For cytokine injections, IL-6 and IFN-γ (Kingfisher Biotech, St. Paul, MN, USA) were prepared as 50 µg/mL stock solutions in PBS and administered intraperitoneally at 10 µL per 0.25 g body weight. TNF-α and CLCF1 (Kingfisher Biotech) were prepared as 250 µg/mL stock solutions in PBS and administered at 10 µL per injection. Human recombinant CXCL11 (Peprotech, NJ, USA) was injected at 10 µL of a 20 µg/mL stock solution.

### EdU labeling and detection

To identify proliferating cells, adult zebrafish were anesthetized in 0.024% tricaine and administered 20 µL of 0.5 mg/mL EdU (5-ethynyl-2′-deoxyuridine; 2.5 mg/mL in PBS) dissolved in HBSS (Sigma Aldrich) via intraperitoneal injection 2 hours before kidneys were harvested. Following the treatment, kidneys were dissected, and EdU incorporation was detected in whole-mount kidney preparations using the Click-iT® Plus EdU Alexa Fluor® 647 Imaging Kit (Thermo Fisher Scientific) according to the manufacturer’s instructions. For dextran filtration experiments, EdU was injected at 6dpi, 10dpi, and 12dpi, and 20 µl 10KD rhodamine dextran (1mg/mL in PBS) (Thermo Fisher) was injected at 8 and 12dpi, and kidneys were harvested at 10dpi and 14dpi.

### In situ hybridization

Whole-mount single in situ hybridization (ISH) was performed as previously described ^85^ with modifications adapted from Kamei et al. ^84^. Digoxigenin (DIG)-labeled antisense RNA probes were generated from zebrafish *lhx1a*, *fzd9b*, and *wnt9b* cDNA sequences ^12^. Adult zebrafish were dissected to remove the head and internal organs, leaving the kidneys attached to the dorsal body wall, and fixed overnight with gentle rocking in 4% paraformaldehyde (PFA; Electron Microscopy Sciences). Following fixation, tissues were washed five times in PBST (1× PBS containing 0.5% Tween-20). Kidneys were dissected using fine forceps and permeabilized with 10 µg/mL proteinase K (Roche) in PBST for 1 h at room temperature with gentle agitation, then post-fixed in 4% PFA overnight at 4°C and washed five times with PBST. Samples were pre-incubated in hybridization buffer (50% formamide, 5× SSC, 50 µg/mL heparin, 500 µg/mL tRNA, 0.1% Tween-20, pH 6.0) at 68°C, followed by hybridization with the DIG-labeled antisense probes at the same temperature. Bound probes were detected using an anti-DIG alkaline phosphatase–conjugated antibody (sheep Fab fragments, 1:5000; Roche) and visualized with the NBT/BCIP chromogenic substrate (Boehringer Mannheim). After staining, kidneys were refixed in 4% PFA, cleared in dimethylformamide, depigmented with hydrogen peroxide, and equilibrated in a 1:1 mixture of PBS and glycerol. Whole-mount samples were imaged using a Leica MZ12 stereomicroscope equipped with a Spot digital camera. For histological analysis, stained kidney tissues were washed in PBT (PBS with 0.5% Tween-20), dehydrated through an ethanol series (50%, 70%, 80%, 90%, 100%; 10 min each), and embedded in JB-4 plastic resin (Electron Microscopy Sciences). Sections (7 µm) were obtained using a Leica RM2165 rotary microtome, mounted with Permount (Fisher Scientific), and imaged using a Nikon E800 microscope equipped with a Spot Insight CCD digital camera.

### Immunohistochemistry

Fish were euthanized, and all organs except the kidneys were removed, leaving the kidneys attached to the body for fixation. Samples were fixed in 4% paraformaldehyde (PFA) for 3 hours at room temperature on a rocker, followed by five washes in PBST (PBS containing 0.1% Tween-20). Kidneys were then dissected, treated with proteinase K for 1 hour, post-fixed in 4% PFA for 1 hour, and washed five times for 20 minutes each in PBST. Whole-mount kidney tissues were incubated overnight at 4 °C in blocking buffer containing the following primary antibodies: chicken anti-GFP (1:5000; Abcam, ab13970) and rabbit anti-DsRed (1:500; Takara Bio, catalog #632496). After primary antibody incubation, tissues were washed in incubation buffer and incubated overnight with secondary antibodies, goat anti-chicken Alexa Fluor 488 (1:3000) and goat anti-rabbit Alexa Fluor 567 (1:300). Tissues were again washed extensively in incubation buffer, followed by nuclear staining with Hoechst 33342 (1:2000) overnight. Finally, samples were washed, mounted in glycerol, and prepared for confocal imaging.

### Kidney tubule isolation

Adult female AB or TuAB trunk kidneys (two per sample) were initially vortexed 30 seconds, then incubated on a rocking stage at room temperature in 5mg/mL Collagenase Type 2 (Worthington) in 0.9X HBSS (calcium, magnesium, no phenol red) (Gibco) for 15 minutes. The tissues were then vortexed for 2 minutes and rocked at room temperature for 15 minutes twice and left standing for 3 minutes to allow tubules to separate from hematopoietic cells by gravity. The supernatant containing the hematopoietic fraction of the kidney was collected and passed through a 40 μm filter (pluriSelect) to exclude any tubules from the preparation before centrifugation at 300xG for 5 minutes. Tubules were washed with 1mL 0.9x HBSS, vortexed, and left standing again for 3 minutes. After discarding supernatant liquid from all samples, tissues were used in an RNAeasy Plus Micro RNA extraction kit (QIAGEN), disrupting the tissue in lysis buffer with a pestle and passing the lysate through a Qiashredder column (QIAGEN) for 2 minutes at 16,000xG. Purified RNA was used in a QuantiTect Reverse Transcription Kit (QIAGEN), yielding cDNA for qPCR analyses in a Lightcycler II (Roche). Depletion of hematopoietic tissue from tubule preparations was confirmed by measuring reduced gata1a expression (qRTPCR) in tubules as compared to the whole trunk kidney.

### RNA sequencing

Raw FASTQ files were processed using the nf-core/rnaseq pipeline (v3.14.0). Ribosomal RNA reads were removed using SortMeRNA (version 4.3.4) prior to alignment. Reads were aligned to the *Danio rerio* reference genome version GRCz11, with annotations from Ensembl release 110 ^86^, using the STAR aligner ^87^ (version 2.79a), and gene-level read counts were quantified using Salmon ^88^ (version 1.10.1) in alignment-based mode. STAR alignment files were manually analyzed for evidence of differential abundance of repetitive reads across samples and conditions at two levels, reads with more than 20 genomic alignments (reported only as a count and fraction of the input reads for each sample), or reads that hit multiply less than 20 times (returned with both alignments and an overall count and fraction of input reads). No significant variation was observed between conditions of either metric. A merged gene counts matrix generated from all sample-specific Salmon runs was used for downstream analysis.

For the early recovery samples, further analysis was performed using R for Mac (version 4.3.3) for mac and using the package DESeq2 ^89^ (version 1.42.1) for correlation of read counts per gene across samples, and test for differentially expressed genes. Poorly expressed genes, those that failed to have at least 10 mapped reads in at least three samples, were removed from the analysis. Read counts of genes for triplicate samples were highly correlated by principal components analysis. DESeq2 was used in “LRT” mode, comparing the model “∼condition” against the base null model “∼1”, reporting genes with significance *padj* <= 0.05. DESeq2’s *rlog* function was used to generate an expression matrix, and then subset to contain only genes that passed the LRT significance test described above. The resulting matrix was Z-normalized gene-by-gene, such that the normalized expression of the *i*^th^ gene in condition j was given by *Z_ij_* = (*r_ij_* - avg_i_)/stdev_i_, where *r_ij_* is the rlog-normalized expression for gene *i*, condition *j* and avg_i_ and stdev_i_, are the average and standard deviation, respectively, for gene *i* across all samples. The resulting Z-matrix was hierarchically clustered with JMP 18.0.1 for Mac, using Ward clustering without column normalization. The final cluster number was selected manually after visual inspection, and gene lists for each cluster used for downstream analysis.

For the 7-day recovery samples, DESeq2 was used with a standard Wald-test to compare expression levels of damaged and control samples, with a single factor design matrix.

### FACS Cell sorting

Adult Tg(cdh17:mCherry) zebrafish kidneys were dissected and processed into single cell suspensions using 0.5% w/v Collagenase Type 2 (Worthington) in 0.9X HBSS (+Ca/+Mg), (Gibco), for 30 minutes for digestion. Collagenase activity was inhibited by the transfer to 0.9X HBSS (-Ca/-Mg) / 2% FBS (Gibco) and were passed through a 40 μm filter (Miltenyi). Cell suspensions were sorted on a BD FACSymphony S6 6-way cell sorter. RNA was isolated using the RNAeasy Plus Micro RNA extraction kit (QIAGEN).

### qRTPCR

RNA was extracted from adult zebrafish kidneys at specified time points after gentamicin-induced injury, or injections of various immunogens – Lipopolysaccharides (LPS), Zymosan, and Poly IC to stimulate inflammation. Extraction was performed using the Qiashredder and RNeasy Plus Universal Kit (Qiagen, Germantown, MD), following the manufacturer’s protocol. RNA was then synthesized into cDNA using QuantiTect Rev. Transcription Kit (Qiagen, Germantown, MD), followed by quantitative PCR (Roche Light Cycler 480) using TB Green-based qPCR kits (Takara) and primers as listed in supplemental Table 6. Each qRTPCR experiment was performed using biological triplicates. Gene expression was normalized to *gapdh* mRNA expression, and data were analyzed using GraphPad Prism.

### Image acquisition

The adult zebrafish kidneys were mounted on glass slides using 1:1 mixture of 50% glycerol and PBS as the mounting medium and a No. 1.5 coverslip (Fisherbrand). Images were acquired using a point scanning confocal microscope (LSM 980, Carl Zeiss Microscopy, Germany) on a Zeiss Axio Examiner Z1 upright microscope stand (ref: 409000-9752-000, Carl Zeiss Microscopy, Germany) equipped with a Plan/Apochromat 10x/0.45 M27(ref:420640-9900-000, Carl Zeiss Microscopy, Germany) and 63x/1.40 Oil objectives (ref: 420782-9900-000, Carl Zeiss Microscopy, Germany). Fluorescence was excited using 488 nm (GFP), 568 nm (mScarlet3), 647 nm (EdU/iFluor 647), and 405nm (Hoechst) lasers, with emission detected using spectral detection windows set to appropriate ranges for each fluorophore. Laser power and detector voltage were adjusted individually for each experiment based on histogram feedback to maintain signal intensities within the linear range, targeting 25–50% of the dynamic range to avoid oversaturation. Images were acquired in confocal mode, 8-bit depth, bidirectional scanning. Z-stacks were collected with both 10× and 63× objectives using optimal Nyquist sampling (1×) in the X, Y, and Z dimensions. Imaging was performed using a motorized scanning stage (130 × 85 PIEZO) with a Z-piezo stage insert (WSB500). The system was controlled using ZEN Blue software (ZEN Pro 3.1, Carl Zeiss Microscopy). Airyscan processing was performed in 3D using automatic settings in super-resolution mode.

### Statistical analysis

Image quantification was performed using Imaris software. Volumetric analysis was applied to z-stack datasets in Imaris to quantify three-dimensional structures of adult zebrafish nephron aggregates. The “Add New Surfaces” feature of Imaris was applied to the green, *lhx1a:eGFP* channel in maximum intensity projections of *lhx1a:eGFP* transgenic kidneys treated with CXCL11. Multicellular, *lhx1a:GFP+* structures with six or more contiguous nuclei and with a volume greater than 1,450 cubic microns were counted as nephron stem cell aggregates. GraphPad Prism 10.1.0 was used to determine *p-*values with one-way ANOVA for more than two groups and *t*-Test for two groups. Significance level is displayed as: not significant (ns) = *p* > 0.05, * = *p* ≤ 0.05, ** = *p* ≤ 0.01, *** = *p* ≤ 0.001.

## Acknowledgements and Funding

This research was supported by the National Institutes of Health (NIDDK 5UC2DK126021; I.A.D., L.O.; R15AI169393 to R.W.), the Harvard Stem Cell institute (DP-0097-11-00; I.A.D.), an NSF REU program grant (NSF DBI 2243416), and NIGMS IDeA Awards under P20GM103423, P20GM104318, and P30GM154610. Heiko Schenk was supported by the German Research Foundation (DFG, SCHE 2173/1-1) and by the PRACTIS - Clinician Scientist Program of Hannover Medical School (MHH), funded by the German Research Foundation (DFG, ME 3696/3-1). Image collection for this manuscript was performed with the assistance of the MDI Biological Laboratory Light Microscopy Facility (RRID:SCR_019166). Animal husbandry was supported by the MDIBL COBRE comparative animal models core (P30GM154610). We thank the cited authors for creating and sharing the transgenic zebrafish used in this study.

## Competing interests

The authors have no competing interests to declare.

## Data and resource availability

GEO/ SRA submission is complete and being processed by NCBI. Acession #’s for transcriptomic data on kidney injury will be supplied prior to publication. *Tg(kim1:mScarlet3)* transgenic zebrafish will be available on request under the standard NIH universal materials transfer agreement for academic research.

## Author Contributions

Conceptualization, O.O., H.S. and I.A.D.; Methodology, C.N.K., H.S., F.B., W.G.B.S., O.A., R.W., L.O. and I.A.D.; Investigation, O.O., C.N.K., H.S., W.G.B.S., O.A., R.U., R.K., E.M., R.C., J.C., R.S., H.F.; Resources, R.W., L.O.; Writing-Original Draft, I.A.D.; Writing – Reviewing and editing, all authors; Supervision, I.A.D.; Funding Acquisition, I.A.D, R.W., L.O., H.S.

## Notes

### Competing Interest Statement

The authors have declared no competing interest.

