## Supplemental figures for "Injury-induced Cxcl11 and neutrophil signaling drive zebrafish kidney regeneration by generating a nephrogenic niche of Fgf and Wnt expression"

**Supplemental figure legends**

**Supplemental figure 1. Macrophage / phagocyte cell counts after acute kidney injury.** (A) Counts of interstitial *kim1:mScarlet3*+ / fluorescent dextran double positive cells per microscope field of control (PBS) and gentamicin injured kidneys. (B) Macrophage marker gene expression (qRTPCR) in control (PBS) and gentamicin injured kidney tissue.

**Supplemental figure 2. Inflammatory gene expression following acute kidney injury.** qRTPCR validation of RNA seq data from acutely injured isolated tubule RNA, 1 and 2 days post injury.

**Supplemental figure 3. Activation of *NFkB:gfp* reporter in tubules that do not overlap with sites of new nephron formation on *slc12a3:mCherry*+ distal tubules.**

**Supplemental figure 4. Dynamics of immunogen-induced inflammatory response vs. gentamicin acute tubule injury induced response.** The immunogens LPS, Zymosan, and PolyI:C induce a rapid cytokine (*il1b, il6, tnfa*) and inflammation marker (*mxa, pim-1*) response in whole kidneys within 3 hours of injection and this response is complete by 24 hours post injection. Gentamicin acute kidney injury does not significantly induce *il1b, il6, or tnfa* at early time points and instead shows delayed induction of the interferon target *mxa* and STAT target gene *pim-1* at 48 and 96 hours post injection.

**Supplemental figure 5. LPS injection does not induce injury marker *Tg(kim1:mScarlet3)* expression in kidney tubules.** Confocal maximum intensity projections show *Tg(kim1:mScarlet3)* expression in scattered interstitial macrophages but not kidney tubules in (A) PBS injected, (B) 2 days post injection of LPS, or (C) 4 days post injection of LPS. Scale bars = 20 µm.

**Supplemental figure 6. Immunogen injection effect on *lhx1a* kidney regeneration marker expression.** LPS injection consistently induced a greater than 10-fold induction of *lhx1a* expression in whole kidney tissue by RTPCR, while zymosan injection was less effective, inducing a greater than 5-fold change in *lhx1a* expression. PolyI:C did not induce *lhx1a* expression, suggesting different TLR pathways are activated by these immunogens.

**Supplemental figure 7. Assaying individual cytokines for kidney regeneration responses.** Intraperitoneal injection of recombinant Clcf1, Il6, Ifn gamma, or Tnfa as described in Materials and methods did not induce a significant change in nephrogenic gene expression (*lhx1a*; qRTPCR).

**Supplemental figure 8. Expression of *cxcl11.1* after acute kidney injury.** Whole mount in situ hybridization of control (A) and 4 days post injury (4 dpi) kidneys (B) shows expression of *cxcl11.1* in patches of interstitial cells (C) adjacent to kidney tubules (T in C). Scale: A, B = 200 µm, C = 20µm

**Supplemental figure 9. Expression of *dusp6* in new nephron aggregates.** Histological sections of wholemount in situ hybridization kidney tissue show that *dusp6* is expressed in similar, new nephron structures in wildtype (A) and *Tg(mpx:mCherry,rac2_D57N)* transgenic kidneys (B), 7 days after gentamicin kidney injury abutting on existing kidney tubules (T). Scale bars = 10μm

### Supplemental figure 1

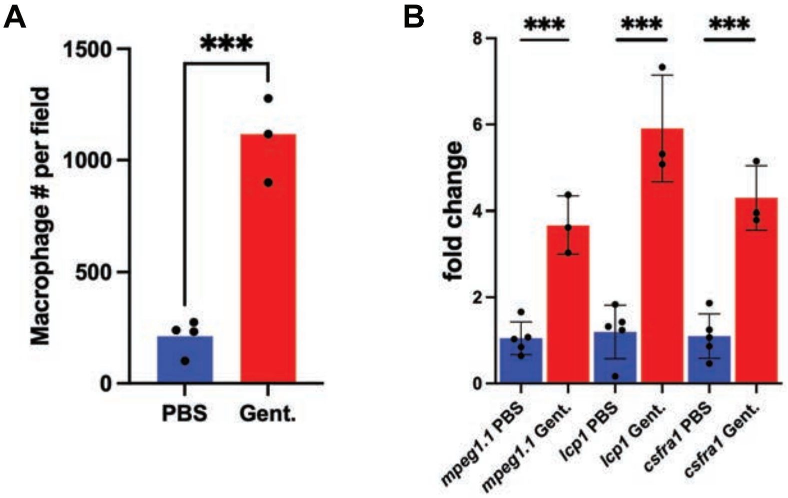

### Supplemental figure 2

● PBS ● Gentamicin

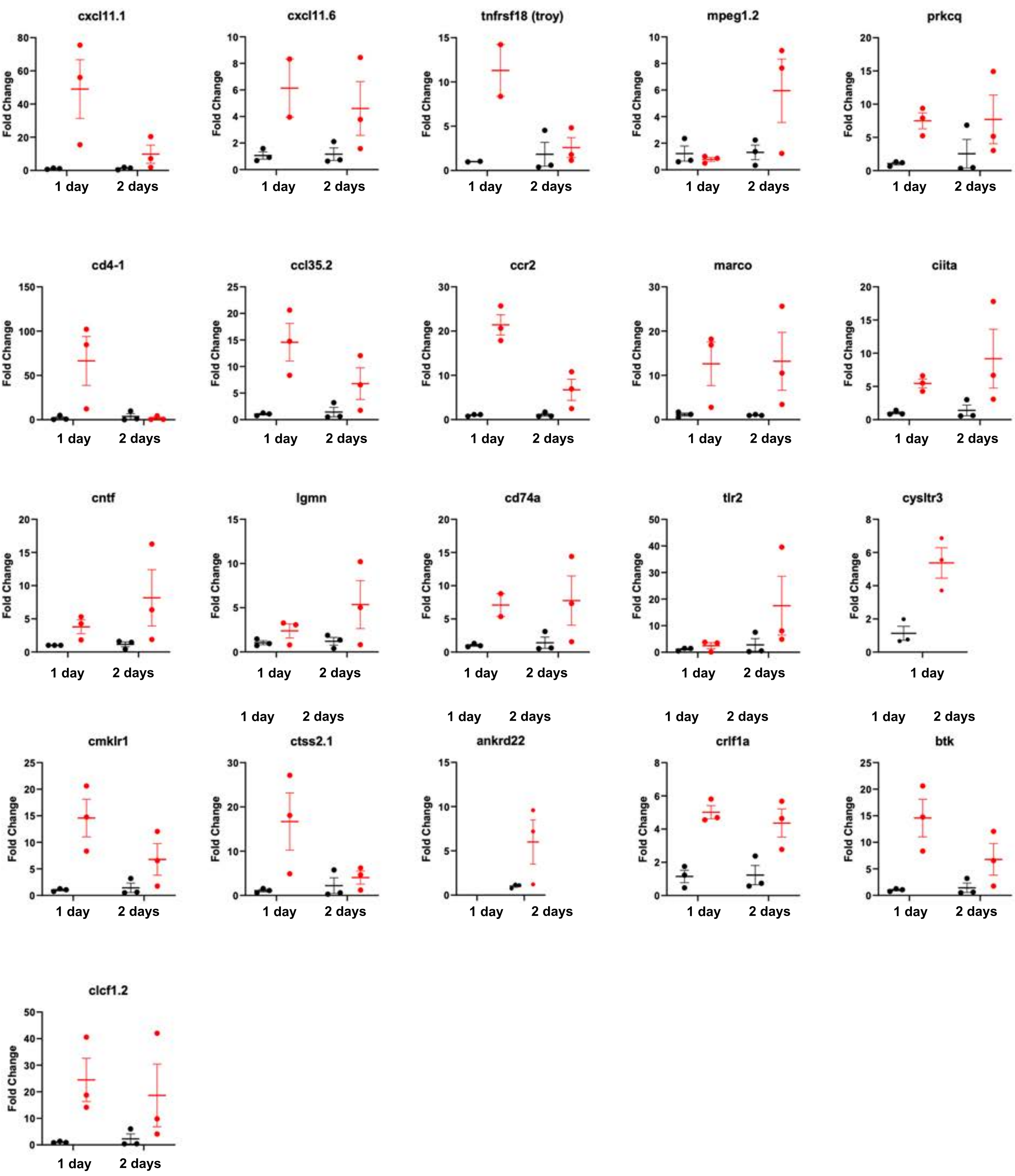

### Supplemental figure 3

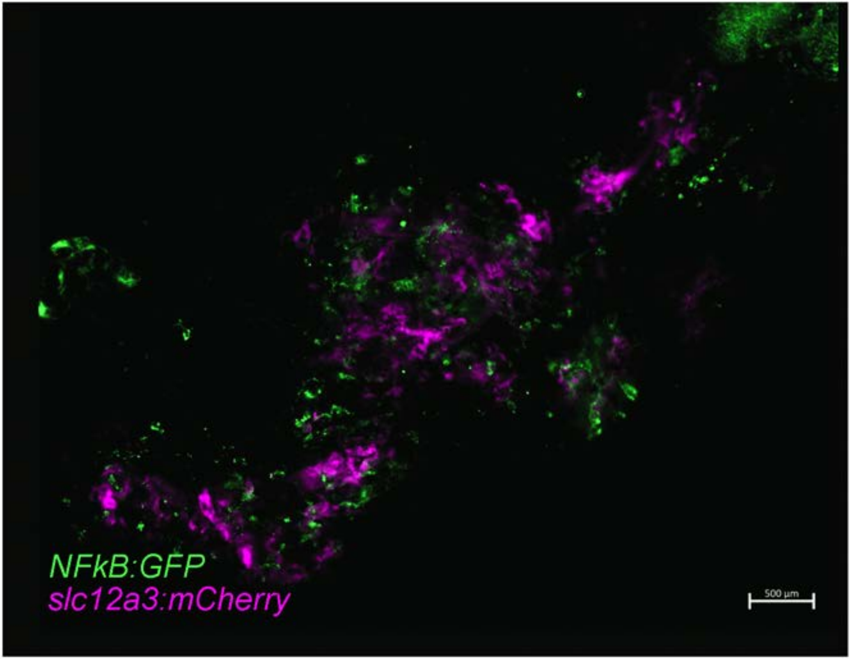

### Supplemental figure 4

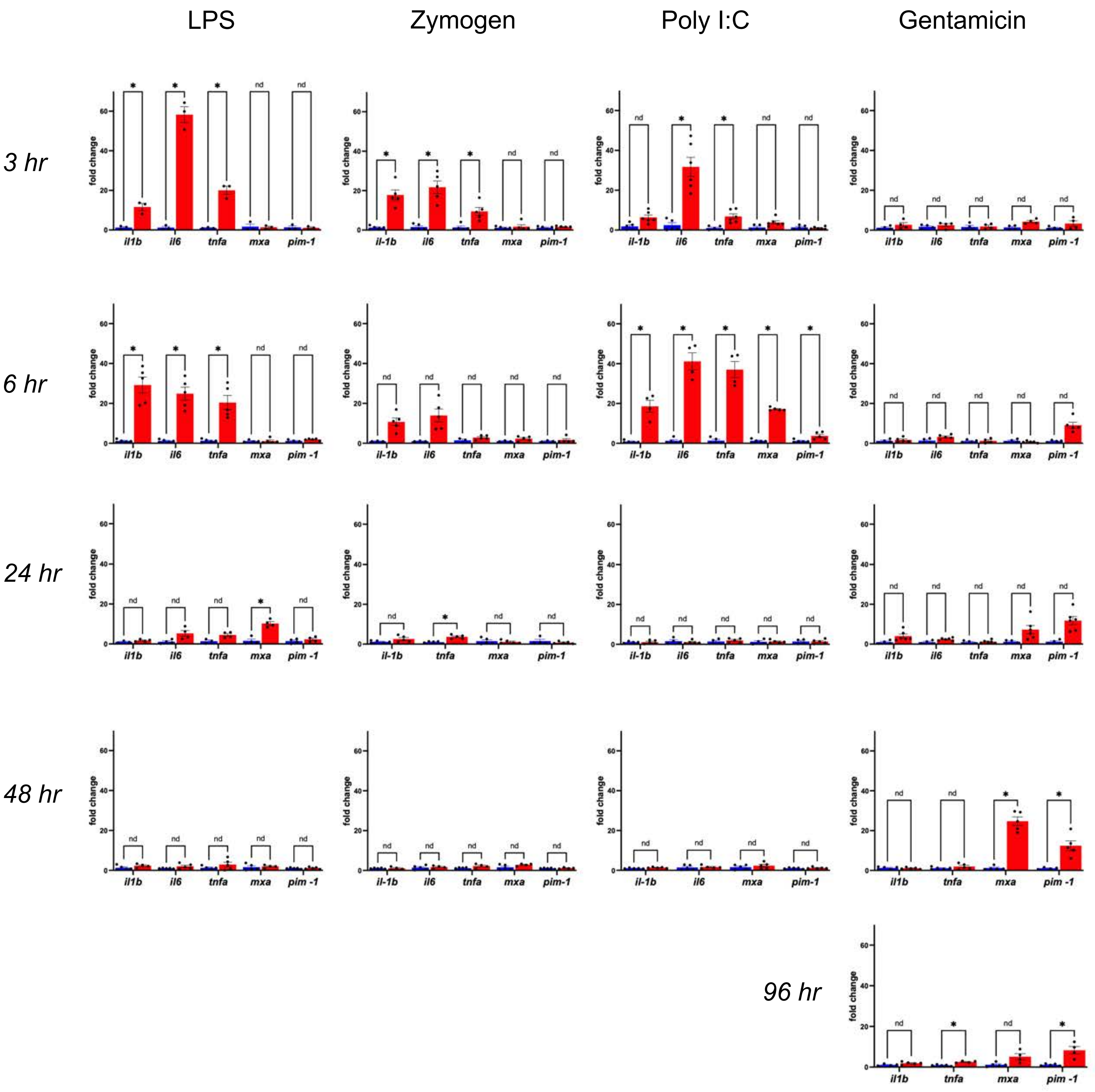

### Supplemental figure 5

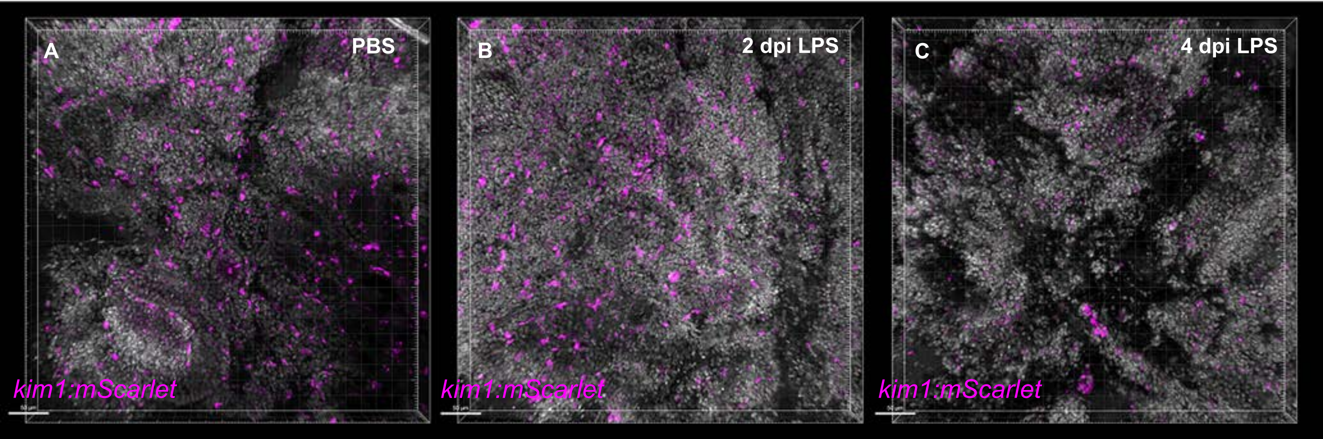

#### Supplemental figure 6

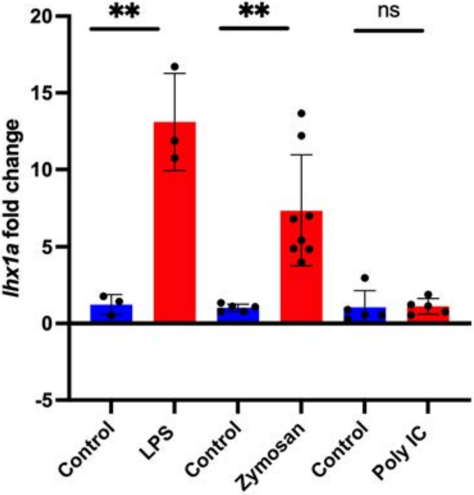

### Supplemental figure 7

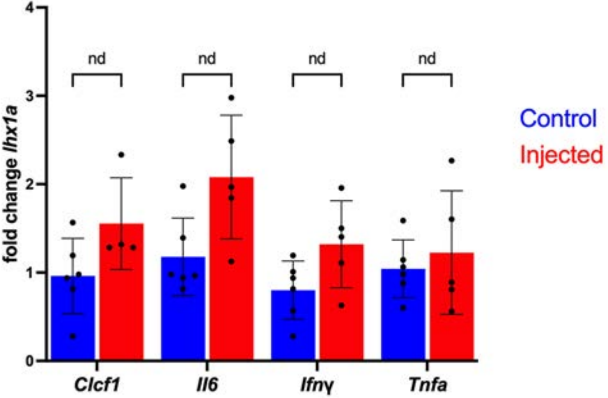

### Supplemental figure 8

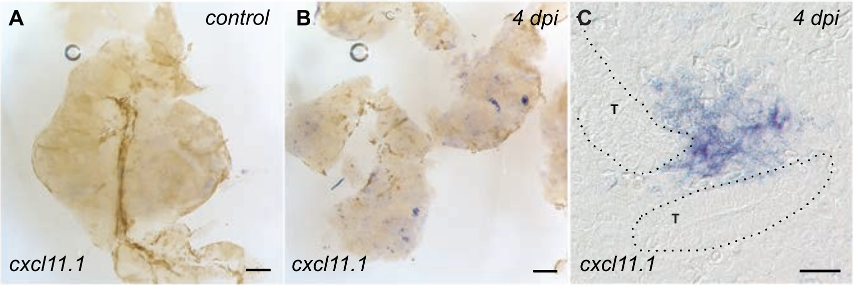

### Supplemental figure 9

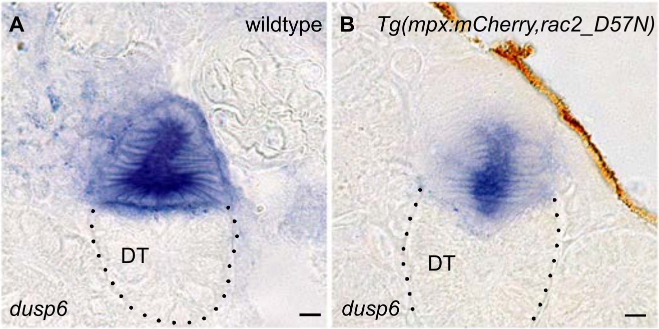
